# *In vitro* ribosome synthesis and evolution through ribosome display

**DOI:** 10.1101/692111

**Authors:** Michael J. Hammerling, Brian R. Fritz, Danielle J. Yoesep, Do Soon Kim, Erik D. Carlson, Michael C. Jewett

**Author notes:** Equal Contributions (Michael J. Hammerling, Brian R. Fritz).

## Abstract

Directed evolution of the ribosome for expanded substrate incorporation and novel functions is challenging because the requirement of cell viability limits the mutations that can be made. However, our recent development of an integrated strategy for the *in vitro* synthesis and assembly of translationally competent ribosomes (iSAT) enables the rapid generation of large libraries of ribosome variants in a cell-free environment. Here we combine the iSAT system with ribosome display to develop a fully *in vitro* methodology for <u>ri</u>bosome <u>s</u>ynthesis and <u>e</u>volution (called RISE). We validate this method by selecting highly active genotypes which are resistant to the antibiotic clindamycin from a library of ribosome variants. We further demonstrate the prevalence of positive epistasis in successful genotypes, highlighting the importance of such interactions in selecting for new function. We anticipate that RISE will facilitate understanding of molecular translation and enable selection of ribosomes with altered properties.

## INTRODUCTION

The ribosome – the macromolecular machine that polymerizes α-amino acids into polypeptides (*e.g.*, proteins) according to messenger RNA (mRNA) templates – is the catalytic workhorse of the translation apparatus. The bacterial ribosome is made up of one small and one large subunit, the 30S and 50S respectively. The 30S subunit is composed of 21 ribosomal proteins (r-proteins) and the 16S ribosomal RNA (rRNA) and is primarily responsible for decoding mRNA. The corresponding 50S subunit is composed of 33 r-proteins, the 23S rRNA, and the 5S rRNA and serves to accommodate tRNA-amino acid monomers, catalyze peptide bond formation, and excrete polypeptides.

The extraordinary synthetic capability of the ribosome has provided the basis for recombinant DNA technology. Recently, engineering mutant ribosomes to elucidate key principles underpinning the process of translation, program cellular function, and generate novel sequence-defined polymers has emerged as a major opportunity in chemical and synthetic biology^1,2^. While efforts to engineer ribosomal function have achieved some success^3,4^, especially with the recent advance of tethered ribosome systems^5,6^, significant repurposing of the translation apparatus is constrained by challenges inherent to altering translation in living cells. A completely *in vitro* approach could overcome cell viability constraints in order to diversify, evolve, and repurpose the ribosome into a re-engineered machine for understanding, harnessing, and expanding the capabilities of the translation apparatus.

In this study, we present a cell-free <u>r</u>ibosome <u>s</u>ynthesis and <u>e</u>volution method called RISE. RISE combines our *in vitro* integrated synthesis, assembly, and translation (iSAT) system with ribosome display^7,8^ (**Fig. 1**). Specifically, a library of rRNA sequence variants is transcribed from plasmid DNA and assembled into a library of ribosomes. Meanwhile, a truncated mRNA encoding a selective peptide is also transcribed in the same reaction. Nascent functional ribosomes within the library then translate the selective peptide and stall to form mRNA-ribosome-peptide “ternary” complexes. These complexes are selectively captured by the peptide, rRNA is isolated and reverse-transcribed, and the resulting ribosomal DNA enters another cycle of RISE for further enrichment of active sequences. In addition to showing that RISE can enrich a target ribosomal genotype from a library by >1000-fold per round of selection, we evolved ribosome resistance to the antibiotic clindamycin. Deep sequencing analysis of this selection revealed a densely connected network of winning genotypes, as well as the first, to our knowledge, experimentally measured estimates of intramolecular epistasis values for a library of ribosome mutants.

**Figure 1.**
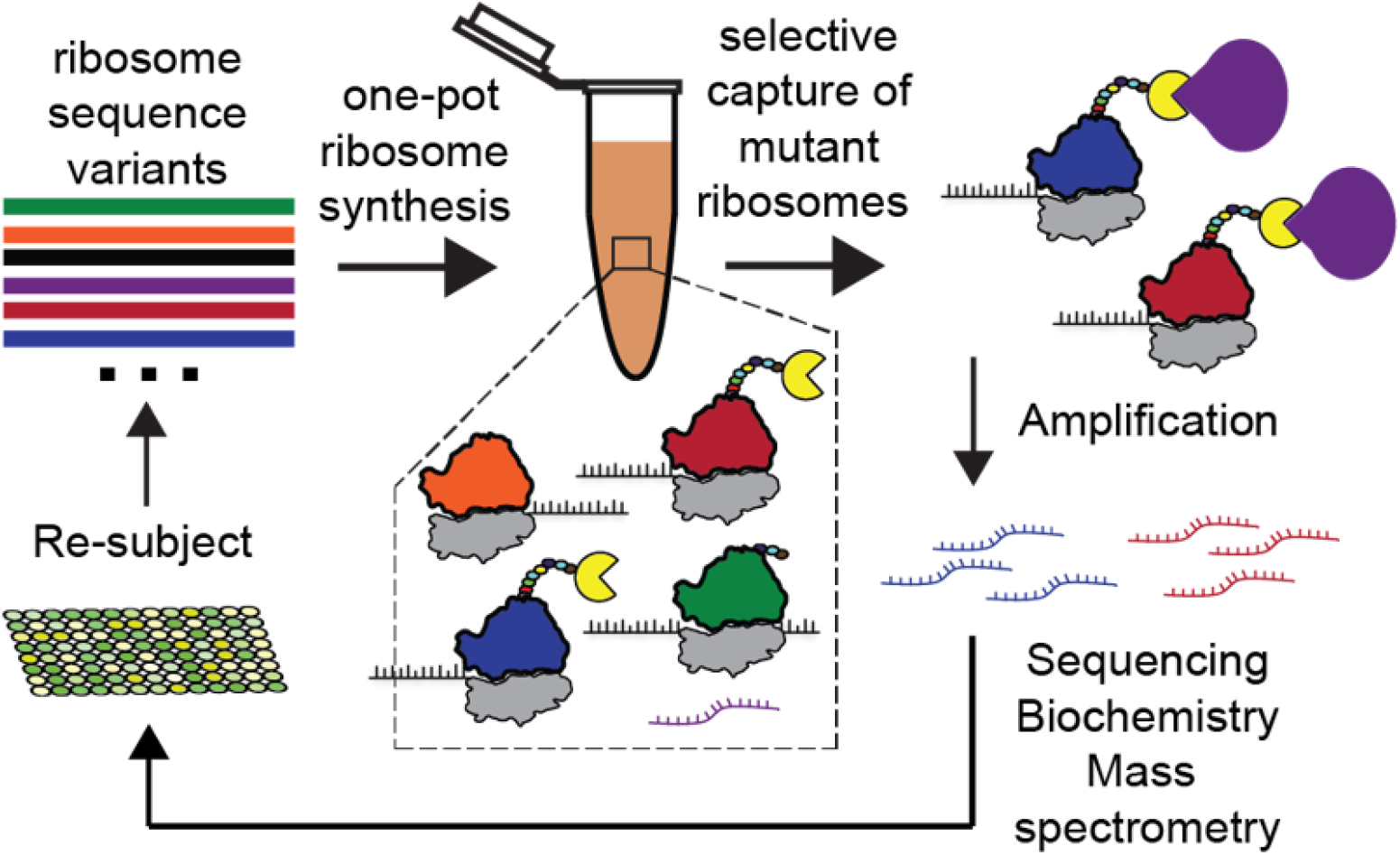
Diagram of *in vitro* ribosome synthesis and evolution (RISE). Starting with a T7-promoted rDNA operon, mutations are introduced to generate an rDNA operon library. The rDNA operon library is then used in an iSAT reaction to assemble a library of ribosomes. Functional ribosomes then translate selective peptides and stall to form ternary complexes, allowing for selection of the peptides and, thus, isolation of functional ribosomes. The rRNA of the functional ribosomes is then purified and reverse transcribed into cDNA for sequencing and reinsertion into the rDNA operon.

## RESULTS

We hypothesized that we could develop RISE by linking iSAT with ribosome display. As a model we focused on *Escherichia coli* ribosomes because the translation apparatus of *E. coli* is the best understood and most characterized both biochemically and genetically^9^. To date, non-physiological conditions and low reconstitution efficiencies from *in vitro* transcribed 23S rRNA have previously represented one of the most serious bottlenecks to constructing and evolving *E. coli* ribosomes *in vitro*^10^. However, we recently addressed these bottlenecks by developing an integrated method for the physiological assembly of *E. coli* ribosomes, called iSAT^9,11–13^. iSAT enables efficient one-step co-activation of rRNA transcription, assembly of transcribed rRNA with native ribosomal proteins into *E. coli* ribosomes, and synthesis of functional protein by these ribosomes in a ribosome-free S150 crude extract. Importantly, iSAT ribosomes possess ~70% of the protein synthesis activity of *in vivo*-assembled *E. coli* ribosomes^11^, which could enable the selection and evolution of ribosomes if the approach were connected to ribosome display.

The development of RISE required the optimization of all aspects of the ribosome display method, including stable ribosome stalling, ribosome capture, and recovery of ribosomal coding DNA (cDNA). We began our investigation by attempting to create stalled mRNA-ribosome-peptide “ternary” complexes. To assess iSAT ribosome stalling, we inserted the gene for superfolder green fluorescent protein (sfGFP) upstream of the spacer sequence in the pRDV vector developed by the Plückthun lab for ribosome display^7,8^. The goal was to gain insight into reaction dynamics and quantify sfGFP production in our iSAT reactions as a proxy for properly stalled ribosomes. Properly stalled iSAT ribosomes should generate one sfGFP molecule for each translating ribosome, with a maximum concentration of 300 nM ribosomes based on the maximal total ribosomal protein concentration in iSAT reactions. Importantly, to ensure that the stop codon of sfGFP did not lead to ribosome release, the pRDV vector was modified to include the self-cleaving hammerhead (HH) ribozyme after the peptide coding region to cleave off the stop codon and generate a truncated mRNA^11,14^. We tested constructs with and without the HH ribozyme in iSAT in the presence and absence of 5 uM anti-ssrA oligonucleotide (**Fig. 2A**). The addition of anti-ssrA oligonucleotides is routinely used in ribosome display to prevent the recycling of stalled ribosomes via the transfer-messenger RNA (tmRNA) encoded by the gene ssrA^15,16^. The combination of the HH gene element and the anti-ssrA oligonucleotide reduced translation to below 300 nM of sfGFP, suggesting that ribosomes are efficiently stalling.

**Figure 2.**
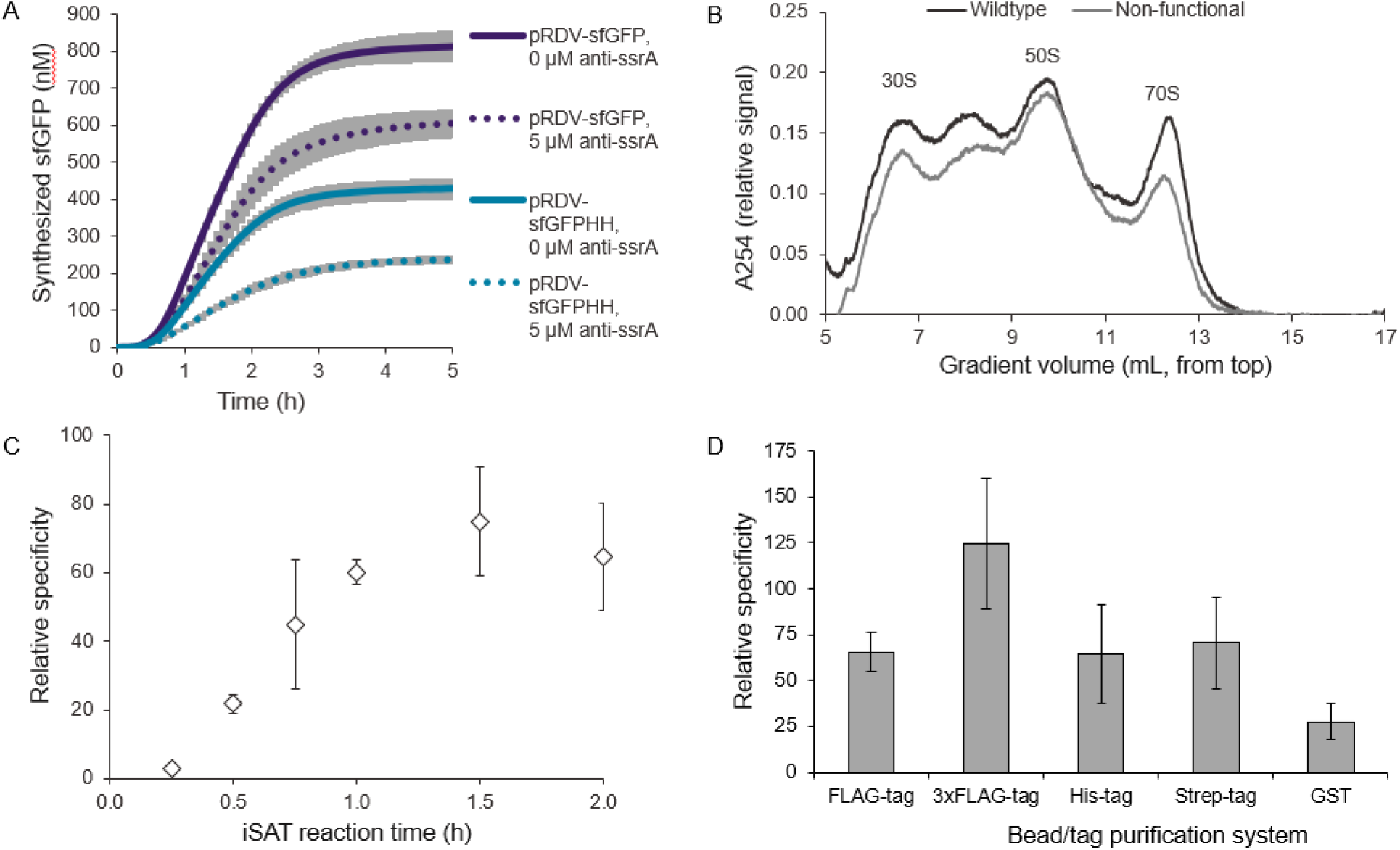
Development and optimization of RISE selection conditions. **(A)** iSAT translation of sfGFP under stalling conditions. pRDV-sfGFP plasmids were made with (blue) or without (purple) a 3’ HH ribozyme for processing of transcribed mRNA to remove the stop codon. Anti-ssrA oligonucleotide was included at either 0 μM (solid) or 5 μM (dotted). **(B)** Sedimentation analysis of iSAT reactions with wild-type or nonfunctional rDNA operon. Peak identities are labeled. **(C)** Relative specificity of capture by anti-FLAG magnetic beads for iSAT reactions displaying FLAG-tag. Reactions were incubated from 15 min to 2 h. **(D)** Comparison of relative specificity of various bead/tag purification systems for use in RISE. Wash methods were held constant for comparison, and reactions were incubated for 1.5 h, except for reactions with GST, which were incubated for 2 h. For **A**, **C** and **D**, values represent averages of three independent reactions (n=3) and shading represents one standard deviation (s.d.).

To assess the specificity of RISE for selecting functional variants, we next carried out RISE with a mock selection whereby reactions contained either wild-type (WT) or nonfunctional rDNA operons that included lethal point mutations in both the 16S and 23S rRNA (ΔC967 and G2252A, respectively)^17,18^. These mutations resulted in an iSAT translation activity of less than 2% of the iSAT activity from WT rDNA operons. From these reactions we determined relative specificity of different RISE conditions by dividing the relative capture of functional ribosomes by the relative capture of nonfunctional ribosomes, as determined by RT-qPCR detection of 23S rRNA in the eluates (**Supplementary Fig. S1)**. Sedimentation analysis supported this approach by demonstrating that nonfunctional rRNA still formed native-like ribosomal particles to preserve potential nonspecific interactions (**Fig. 2B**).

With the observations of efficient ribosome stalling from actively translating ribosomes, we set out to optimize the reaction duration, which has been shown to be critical for optimizing capture of stable ternary complexes^19^. We observed that iSAT reactions require approximately 30 minutes for rRNA synthesis, ribosome assembly, and translation of detectable levels of sfGFP (**Fig. 2A**). By varying iSAT reaction times, we observed that relative RISE specificity is highest at 1.5 hours (**Fig. 2C**). Visualization of the recovered nucleic acids from 1.5-hour reactions shows that reactions with nonfunctional rDNA operon plasmids do not show visible rRNA capture, whereas reactions with functional WT rDNA operon plasmids show bands representative of 23S and 16S rRNAs (**Supplementary Fig. S2**).

The key conceptual shift in RISE from conventional ribosome display is that rRNA sequences are selected rather than mRNA sequences. Since ribosome mutants are the focus, we next explored conditions for optimal ribosome capture. The idea was to synthesize an N-terminal peptide handle (*e.g.*, the FLAG-tag) to emerge first from the ribosome and be displayed for affinity purification and capture. Several common selective peptide tags were tested, including the FLAG-tag, 3xFLAG-tag, His-tag, Strep-tag, and glutathione S-transferase (GST), in the pRDV vector using commercially available magnetic capture beads. iSAT reactions expressing each tag or protein were incubated for 1.5 h, except for GST, which was incubated for 2 h to account for the additional translation and folding times associated with expressing a large selective protein. Given the results of this screen, we elected to use the 3xFLAG-tag peptide in RISE reactions, which exhibited a 124-fold specificity for functional over inactive ribosomes (**Fig. 2D**).

We next optimized binding and wash conditions for the captured ternary complex displaying the 3xFLAG-tag. Addition of bovine serum albumin (BSA), Tween® 20, or heparin to the binding and/or wash buffers were tested for their ability to decrease nonspecific binding between the magnetic bead and the 3xFLAG-tag. BSA and Tween® 20 were found to increase relative specificity for active ribosomes by 25% and 19% respectively, while heparin had the counterintuitive effect of decreasing relative specificity (**Supplementary Figures S3A and S3B, Supplementary Table S3**). Most importantly, we found that RISE specificity was greatly improved by increasing the stringency of the washing steps, with increasing the number of wash steps from 5 to 10 providing the greatest improvement (**Supplementary Figures S3C and S3D**). While we observed that specific capture is unchanged by increased wash stringency, nonspecific capture was significantly lowered, resulting in increased relative specificity of >1,000-fold per round of selection (**Supplementary Figure S3D**).

To complete RISE, we developed a strategy to reverse-transcribe recovered rRNA sequences and reinsert the recovered cDNA into the operon plasmid. Given constraints in reverse transcribing full-length rRNA, which may contain post-transcriptional modifications, initial focus was given to manipulation of only the peptidyl transferase center encoded by the 23S rRNA. We used RT-PCR to recover a 660 bp region of the 23S rRNA gene (bases 1962 to 2575, *E. coli* numbering), and off-site type IIs restriction digestion and ligation to insert the 660 bp fragment from the 23S rRNA gene into pT7rrnBΔ660, an rDNA operon plasmid lacking the amplified gene fragment. Assembled libraries were transformed into cells to maintain a stable plasmid stock, and this plasmid pool was used in the next round of *in vitro* selection.

With the RISE platform at hand, we next sought to use it to select and evolve mutant ribosomes. We chose resistance to the ribosome-binding antibiotic clindamycin as a model system^20^. Previously, Cochella and Green demonstrated that ribosome display can be used for the selection of functional ribosomes from a ribosome library, but their method required *in vivo* ribosome synthesis and ribosome purification, which biases and reduces ribosome library size as well as limits throughput^20^. RISE avoids these requirements, which should facilitate the development of rapid, high-throughput selection strategies for ribosome evolution. We developed a clindamycin-resistance (CR) library by randomizing positions 2057 to 2062 of the 23S rRNA, a region associated with clindamycin binding^21^. We then applied RISE to the CR library in the presence of 500 μM clindamycin in iSAT reactions. As a control, we carried out selections in the absence of clindamycin to evaluate the impact of selection pressure on the RISE method. Before selection, the libraries demonstrated no detectable protein synthesis due to the prevalence of inactive ribosomal variants. After four rounds of selection, the library selected in the absence of clindamycin (Clin−) demonstrated 1.5-fold greater bulk-protein synthesis than the library selected in the presence of clindamycin (Clin+) (**Fig. 3A**, green bars). Conversely, the Clin+ library improved to have 8.7-fold greater bulk protein synthesis activity after the completion of four RISE cycles in the presence of clindamycin than the Clin− library (**Fig. 3A**, purple bars). These results are commensurate with the conditions under which the libraries were selected and demonstrate the ability of RISE to select ribosomal genotypes that are tailored to functioning optimally under a given selective challenge.

**Figure 3.**
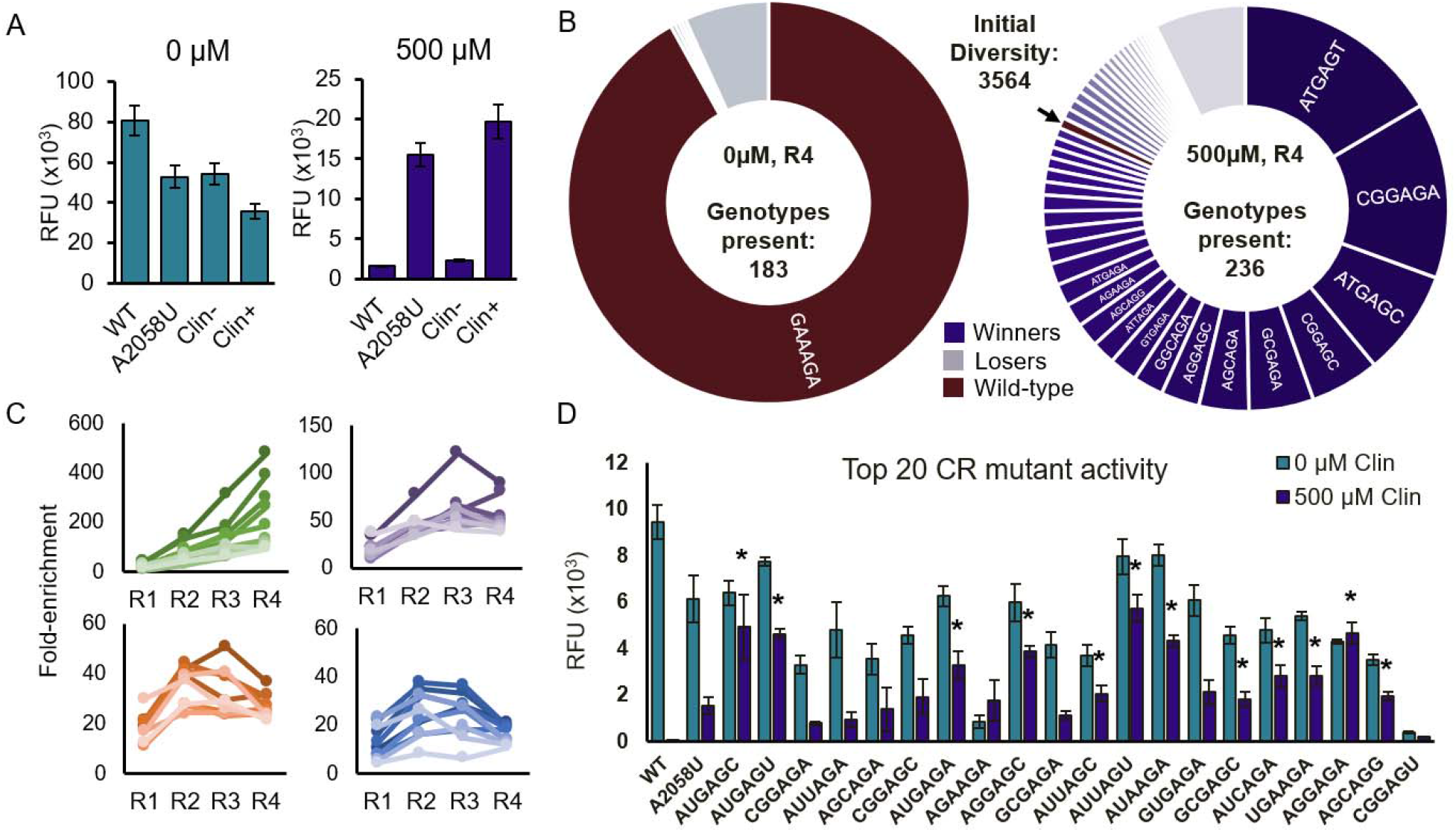
Results of selection of CR ribosomes in RISE. After four rounds of selection in the absence (0 μM) or presence (500 μM) of clindamycin, the resulting selected libraries were analyzed. **(A)** In 0 μM clindamycin (green bars), WT ribosomes maintain the highest expression of sfGFP in iSAT. The A2058U mutant and Clin− selection pool also have high activity, while the Clin+ selection pool has the lowest activity, though all samples express sfGFP. In contrast, in 500 μM clindamycin (purple bars), the WT and Clin− conditions show almost no sfGFP expression, while the A2058U mutant and Clin+ selection pool retain robust expression. Notably, the results of the Clin+ selection pool have a higher activity than the A2058U mutant, suggesting that this pool contains mutants with higher activity in the presence of clindamycin. **(B)** Donut plot depicting the frequency of the top mutants ranked by final frequency after four rounds of selection in 0 μM (left) and 500 μM (center) clindamycin. While WT has nearly taken over the population after four rounds of selection in the 0 μM case, the 500 μM case retains much genotypic diversity, and WT is being rapidly supplanted by CR genotypes. WT is indicated with an arrow on the 500 μM plot. **(C)** Selection dynamics of top genotypes 1-10 (green), 11-20 (purple), 21-30 (orange) and 31-40 (blue) in the 500 μM selection as ranked by final fold-enrichment. Selection dynamics are consistent with a constant selection pressure where very-fit genotypes increase in frequency throughout the course of the selection, while moderately fit genotypes increase initially, but begin to decrease as the average fitness of the population surpasses their fitness. **(D)** The activity of WT, A2058U, and the top 20 genotypes from the Clin+ selection as ranked by fold-enrichment at the end of the selection (n=7). All but one mutant is active in iSAT, and the majority (12 of 20) have higher activity in the presence of clindamycin than the A2058U mutant discovered in other studies (* indicates *p*-value < 0.05).

To achieve high resolution analysis of selection dynamics, we performed deep sequencing of the initial CR library and at each round of selection. Using our quality control criteria (**Materials and Methods**), 3564 total genotypes of 4096 possible genotypes were observed in the initial library (**Supplementary Fig. S4)**. The 0 μM selection resulted in a rapid convergence of the library towards the WT 23S rRNA sequence, though 183 genotypes were still present by the end of the selection (**Supplementary Fig. S5**). In contrast, the 500 μM selection resulted in the depletion of WT 23S rRNA sequence and a more heterogeneous overall pool which converged to 236 genotypes by the final round of selection. From four rounds of RISE on the CR library, the ten most abundant of these genotypes (which comprised 0.38% of the initial library), made up ~64% of the final population, indicating rapid convergence to a handful of the most active CR genotypes (**Fig. 3B**, **Supplementary Fig. S6**). Selection dynamics of the top 40 genotypes show the expected trends of a consistent selection pressure, with the fittest genotypes increasing in frequency throughout the course of the selection (top left panel) while moderately fit genotypes increase initially, then decrease in frequency in later rounds as the average fitness in the population increases (**Fig. 3C**). To re-assess optimal RISE incubation time, but now with finer resolution than before (**Fig. 2C**), we performed time course RISE reactions in which CR genotypes were enriched from degenerate library in 500 μM clindamycin. We then compared enrichment of each genotype after four rounds of selection against its enrichment when the first round of selection is harvested at time points from 7.5 to 90 minutes (**Supplementary Fig. S7**). Supporting our initial bulk data, deep sequencing analysis confirmed efficient selection occurs for 90 minute time points.

A simple method for calculating expected final frequency of a genotype in the population that ignores interactions between positions entails multiplying the final frequency at each selected position together to obtain an expected final frequency for the genotype. Many winning sequences differ substantially from their expected frequency. For example, the consensus “winner” AUGAGA sequence is at only 33% the expected final frequency based on this method, though it is still present in the top 10 sequences by fold-change. In contrast, two other top sequences, GCGAGC and UGAAGA were found at 10.1-fold and 17.8-fold higher frequency than expected (**Supplementary Figure S8**). These unexpected winners highlight the importance of creating diverse libraries that sample broad sequence spaces to discover difficult-to-predict sequences that may have high activity in the given selective challenge.

After completion of the *in vitro* evolution of the CR library, the top 20 genotypes from the 500 μM condition (ranked by their fold-enrichment at the end of the selection) were individually cloned and tested in iSAT. All genotypes translated sfGFP in the presence of 500 μM clindamycin, indicating successful selection for clindamycin resistance (**Fig. 3D**). A majority (12 of 20) were more active in iSAT reactions with clindamycin compared to the known A2058U mutation previously demonstrated to confer clindamycin resistance *in vivo* and also shown in **Figure 3D**. Surprisingly, none of our most active CR mutants were observed previously in *in vitro* clindamycin-resistance selections^20^.

To understand why some of these sequence variants had not been previously observed, we assessed whether the RISE platform discovered variants that were non-viable *in vivo*. To do so, we exchanged the T7 promoters of the top 20 CR operon variants with *P*_*L*_ promoters for *in vivo* transcription by native polymerases. These constructs were then transformed into the Squires strain, which lacks rRNA sequences on the genome and instead derives rRNA from a plasmid-based operon. The Squires strain serves as a testing platform for whether an rRNA sequence (encoded by the plasmid) can support life, and generally if such rRNA sequences are compatible with cell growth^22^. Transformants were plated on: i) carbenicillin, ii) carbenicillin and sucrose, or iii) carbenicillin, sucrose, and clindamycin, to assess the ability of these ribosomes to support life (**Fig. 4A, B**). Carbenicillin selects for the presence of our variant rDNA operon plasmid, sucrose counterselects the WT rDNA already present in the Squires strain (through the SacB negative selection marker), and clindamycin tests if the clindamycin-resistance phenotype also transfers to *in vivo* settings. The CR variants showed no inherent toxicity to the cells when plated on carbenicillin alone, though only 8 of 20 variants were clearly capable of supporting cell growth upon loss of the WT rDNA operon on sucrose and clindamycin agar, suggesting that the other variants do not form viable ribosomes *in vivo*. However, two mutants (CR3, CR5) inviable on plates supported cell growth in liquid culture (**Fig. 4C)**. Of the CR mutants that were viable *in vivo*, there was no clear relationship between fold-enrichment after RISE and cellular growth rate. Interestingly, the non-viable variants all contained mutations at the 2062 position, which is consistent with the previous observation that such mutations are lethal despite *in vitro* ribosome activity^20,23^. The result that ribosomes that are highly active *in vitro* may be inviable *in vivo* demonstrates the potential of RISE for selecting ribosomes that are proficient at translating challenging templates or monomers without concern for *in vivo* viability.

**Figure 4.**
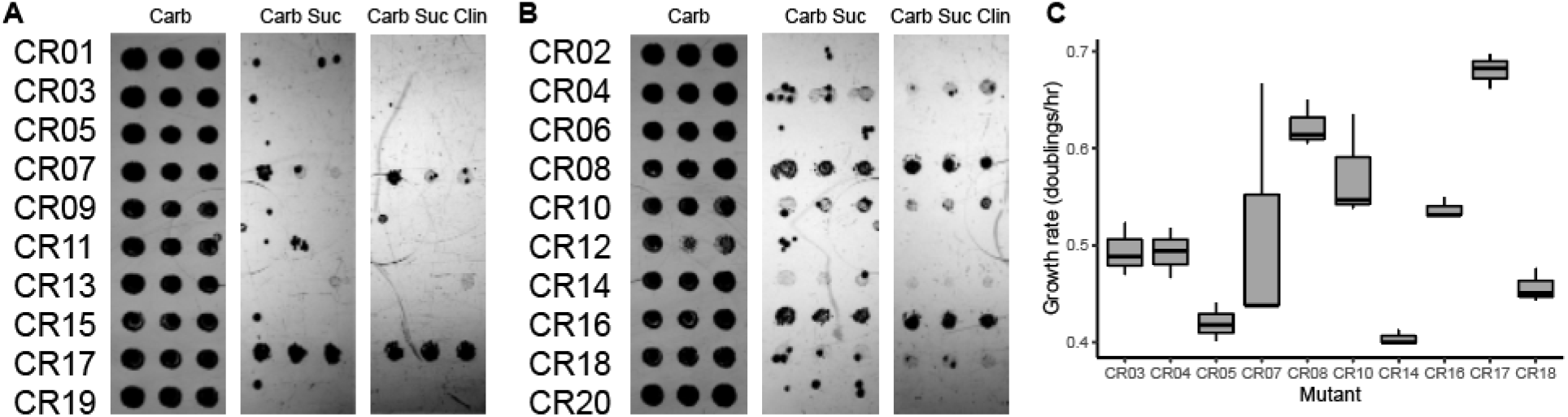
Viability of CR ribosome variants *in vivo*. Isolated Clindamycin Resistant (CR) rDNA variants were altered to include a native promoter for *in vivo* expression. **(A,B)** The top 20 variants were transformed into the Squires strain, and cells were spotted on (left) LB with 100 μg/mL carbenicillin, (center) LB with 100 μg/mL carbenicillin and 5% w/v sucrose, and (right) LB with 100 μg/mL carbenicillin, 5% w/v sucrose, and 350 μg/ml clindamycin. Cells are expected to grow in the first column unless the ribosomes are dominant-lethal, and all spots grow as expected. In the second column, the WT ribosomal operon should be lost, and cells should only grow if the mutant ribosome can support life or if successful escape mutants evolve. Finally, in the far-right column, escape mutants are unlikely due to the additional constraint of clindamycin resistance, and spots should only grow if the CR mutant provided can support life. Mutants CR04, CR07, CR08, CR10, CR14, CR16, CR17, and CR18 are all observed to support life on plates. **(C)** The mutants were also grown in LB carbenicillin sucrose clindamycin liquid cultures (n=3). An additional two mutants, CR03 and CR05, support cell growth in this liquid medium compared to solid medium, though with a very low growth rate. These mutants are arranged in decreasing order of final fold-enrichment in the RISE selection from left to right. Notably, final fold-enrichment does not predict growth rate *in vivo*, supporting the notion that RISE can select ribosome variants that would not be favored in *in vivo* selections.

We next sought to understand the role that epistatic interactions played in our most active genotypes. Epistasis – the nonadditive impact of combinations of mutations within or between genes – is broadly believed to play an important role in the evolution of biomolecules and biological systems^24^. Despite this, little experimental work has been done to measure the degree to which epistatic interactions impact ribosome function^25^. The combination of RISE with next generation sequencing analysis allowed us to estimate epistasis values of genotypes for examining general trends and identifying unexpected functional sequences. We calculated an expected enrichment for each genotype assuming an independent contribution of each nucleotide to the fitness of CR mutants (**Materials and Methods**).

To gain both an intuitive and a quantitative understanding of the relationship between fitness, epistasis, and relatedness in our CR fitness landscape, we visualized all surviving CR mutants as a network plot of point-mutant-adjacent genotypes (**Fig. 5A**). This network has several salient features that suggest that RISE is an efficient selection strategy that enriches genotypes in a manner commensurate with their fitness in the selection. If the impact of each individual residue on overall ribosome fitness is independent, the actual enrichment of each mutant will correlate closely to this expected enrichment, and the log-transformed epistasis values will be close to zero. The distribution of values for those genotypes for which epistasis could be calculated is approximately normally distributed (Shapiro-Wilk test, *p* = .117) with a mean slightly above zero (one-sided Student’s t-test, p = .005) (**Fig. 5B**, gray bars). This slight positive bias may result from the fact that epistasis values could not be calculated for genotypes which dropped out of the population after one round of selection. Average log-fold enrichment is similar for genotypes 2-4 mutations away from wild-type (*i.e*., Hamming distance), though the 7 fittest variants all have 3-4 differences (**Fig. 5D**). Average epistasis increases with greater distance from wild-type up to a Hamming distance of 4 (**Fig. 5E**). Mutants with a Hamming distance of 5 do not conform to the general trend on either plot, likely because genotypes containing 5 mutations necessarily contain a mutation at positions 2060 or 2061, which are important for translation function, and are nearly completely converged to wild-type in our selection (**Supplementary Figure S8**).

**Figure 5.**
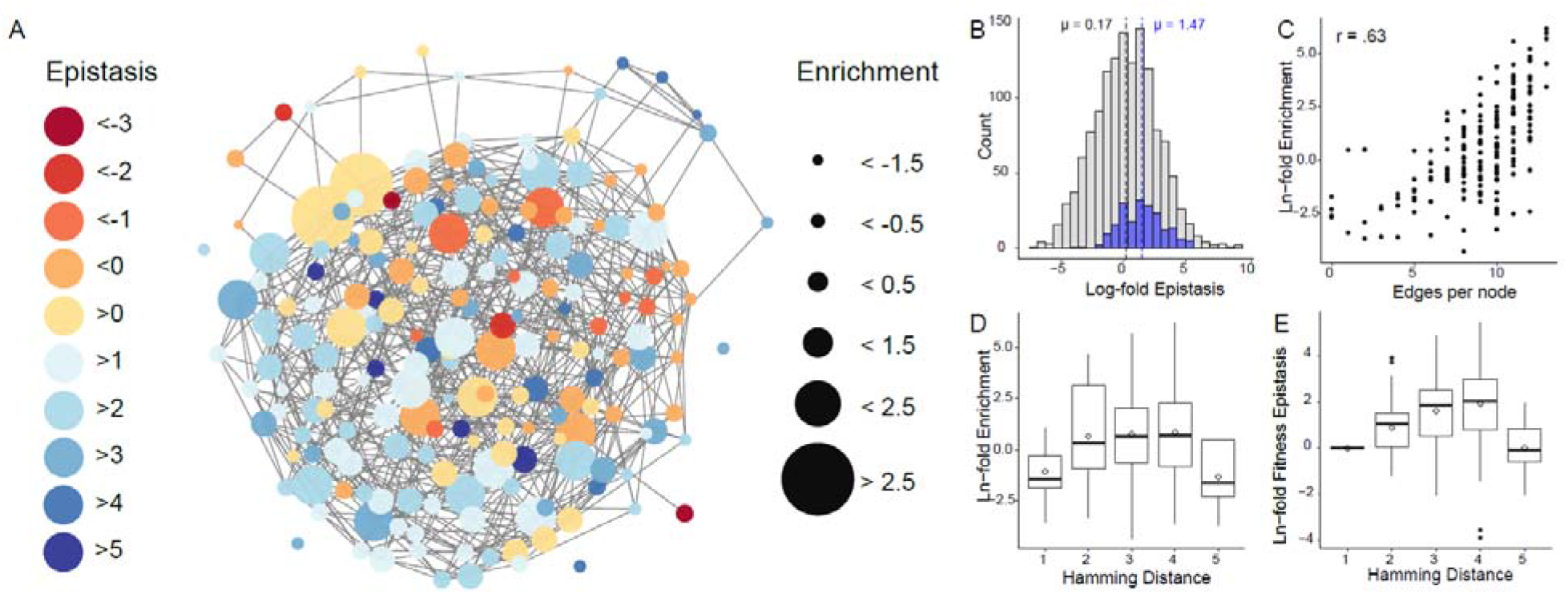
Quantitative analysis of fitness and epistasis of the network of top Clindamycin Resistant genotypes. **(A)** Network plot of Clindamycin Resistant genotypes from RISE selection. In this network plot, each node represents a surviving genotype after four rounds of selection in 500 μM clindamycin. Edges connect nodes that are a single point mutation away from each other. The color of the node represents the calculated epistasis value for that genotype. The size of the node represents the log_10_ of the final enrichment at the end of the selection. This transformation was performed to enable depiction in the network plot on a scale appropriate for publication. This network plot may be interpreted as a visual representation of the combined fitness and epistasis landscapes of the top CR genotypes. **(B)** Average epistasis value of winning genotypes is substantially more positive (μ = 1.47, p = 2.2 × 10^−16^, one-sided Student’s t-test) compared to the overall population of mutants that survived after one round of selection (μ = 0.17). **(C)** Relatedness to other winning genotypes in the population (Edges per node) is correlated with enrichment. **(D)** Box plots of CR mutants that are 1-5 mutations away from wild-type (Hamming distance) are depicted with the natural log transformation of final fold enrichment after selection in clindamycin or **(E)** epistasis values. Both values increase with distance from wild-type except in the case of Hamming distance = 5 mutants, which necessarily entail mutations to highly conserved nucleotide positions.

Edges per node is a measure of the relatedness of a winning genotype to other winning genotypes in the final CR population. This measure correlates well with enrichment (Pearson’s r= .63), indicating that genotypes that are closely related to other genotypes in the final population are most likely to have high fitness (**Fig. 5C**). The high degree of relatedness between the most enriched genotypes is encouraging, as it implies that RISE efficiently selects on some shared property of these closely related genotypes – presumably translational fitness. This is backed up by the high measured translational activity of these genotypes (**Fig. 3D**). In contrast, edges per node does not correlate with epistasis (Pearson’s r = −.09) – indicating that relatedness of genotypes is not predictive of their propensity for epistatic interactions (**Supplementary Figure S9**). Finally, we observed a highly significant increase (one-sided Student’s t-test, *p* = 2.2 × 10^−16^) in average epistasis values in the winning CR genotypes compared to the overall population, indicating that positive epistasis plays an important role in determining the fittest genotypes in this selection (**Fig. 5B**). Future experiments with other selections will determine whether the importance of positive epistasis in the ribosome is a general phenomenon, or if each selection experiment produces different epistatic trends.

To test the function of some of the most unusual top positive epistasis mutants, we cloned 8 of the mutants with variation at positions 2060 or 2061 – the most converged positions in our library, which are generally thought to require wild-type nucleotides for function^26^. Several of these mutants were functional in iSAT, demonstrating that their measured positive epistasis was a real effect, and wild-type nucleotides at these positions are not strictly required for translation (**Supplementary Fig. S10**). These results in epistatic interactions in the ribosome suggest the importance of deep epistasis scanning experiments to acquire a high-resolution map of the positional interactions that govern ribosome function.

## DISCUSSION

In this study, we present the RISE method as a tool for evolving ribosomes *in vitro*. This was accomplished by uniting *in vitro* ribosome synthesis, assembly, and translation with ribosome display. By optimizing reaction conditions, we show that RISE can enrich a target 23S rRNA gene from a library >1000-fold per round of selection. We applied RISE to the *in vitro* evolution of ribosomes from a complex pool of variants, rapidly enriching for clindamycin resistance across four rounds of selection. The inability of some recovered rDNA variants to support cell growth shows that RISE can access genotypes that would not be recoverable from an *in vivo* system, allowing us to explore more diverse functional sequence space of the translation machinery.

Looking forward, we anticipate that RISE will serve as a valuable tool to broaden efforts to expand the range of genetically encoded chemistry into proteins. This is because RISE removes the restrictions of cell viability, transformation efficiency, and growth delays encountered by *in vivo* studies. With this new approach, ribosomal mutants can be rapidly created and screened, and in conditions that would not be possible in cells (*e.g*., non-natural pH or redox environment, or in the presence of additives that are toxic to cells). RISE also holds promise to map the mutability and epistatic interactions of the *E. coli* ribosome, as well as dissect *in vitro* ribosome assembly and function in fine detail. Based on the prevalence of positive epistatic interactions in our winning CR genotypes, we predict that such interactions will be critical to engineering new function in the ribosome. We also posit that RISE will facilitate efforts to both understand the fundamental constraints of the ribosome’s RNA-based active site and create biopolymers containing mirror-image (D-α-) and backbone-extended (β- and γ-) amino acids^27–29^. Such polymers could accelerate the discovery of next-generation therapeutics and materials^30^.

## MATERIALS AND METHODS

### Plasmid and library construction

The plasmids pT7rrnB (containing rDNA operon *rrnB*) and the reporter plasmids pK7LUC and pY71sfGFP were used in iSAT reactions as previously described^10,11^. A variant of pT7rrnB with a 660 bp deletion in the 23S gene, named pT7rrnBΔ660, was created by inverse PCR as previously described^3^ (**Supplementary Table S1**).

Plasmids for ribosome display were constructed from the pRDV plasmid^7,8^. For selective peptide or protein gene insertion, the gene was first amplified by PCR with primers encoding a 5’-GGTGGT-3’ spacer and restriction sites for either NcoI for forward primers or BamHI for reverse primers (**Supplementary Table S1**). The amplified genes and pRDV were digested with NcoI and BamHI, and the correct fragments were isolated by gel electrophoresis and extracted. Fragments were ligated with Quick Ligase (NEB) and transformed into chemically competent DH5α cells, plated, and grown overnight. Resulting isolated colonies were grown for plasmid purification and sequencing.

Libraries of rDNA operons were created from the pT7rrnB plasmid through PCR amplification of particular rDNA fragments with phosphorylated primers containing overhangs of degenerate bases (**Supplementary Table S1**). DNA fragments were ligated and PCR amplified for *in vitro* insertion into the pT7rrnB plasmid (see below).

### iSAT reactions

iSAT reactions were performed as previously described^11,12^. Briefly, salts, substrates, and cofactors were mixed with 1 to 4 nM reporter plasmid and a molar equivalent of pT7rrnB. For ribosome display reactions, the anti-ssrA oligonucleotide (5’-TTA AGC TGC TAA AGC GTA GTT TTC GTC GTT TGC GAC TA-3’) was included at 5 μM to prevent dissociation of stalled ribosomes. Then a mix of proteins was added to final concentrations of approximately 2 mg/mL S150 extract, 300 nM total protein of the 70S ribosome (TP70), and 60 μg/mL T7 RNA polymerase. Reactions were mixed gently by pipetting and incubated at 37°C. Replicate reactions to test translational activity of clones were assembled using the Echo 525 acoustic liquid handling robot to minimize error in pipetting. Preparation of S150 extract, TP70, and T7 RNA polymerase were performed as previously described^10–12^. For sfGFP production, quantification was performed as previously described^10^.

### Sedimentation Analysis

Sedimentation analysis was performed as previously described^12^. Briefly, ribosome profiles were determined from 50 μL iSAT reactions by incubating reactions for 2 h at 37°C, loading them onto a 10-40% sucrose gradient made with Buffer C (10 mM Tris-OAc (pH = 7.5 at 4°C), 60 mM NH4Cl, 7.5 mM Mg(OAc)2, 0.5 mM EDTA, 2 mM DTT) and ultra-centrifuging in SW32.1 tubes at 35,000 rpm and 4°C for 18 h. Gradients were then analyzed through spectrophotometry and fractionation (500 μL fractions). Ribosome profiles were generated from absorbance of the gradient at 254 nm and peaks were determined from comparison to previous traces^12^.

### Ribosome Synthesis and Evolution (RISE)

Fifteen μL iSAT reactions were performed as described^12^, using pRDV_3xFLAG as the ribosome display template and with the addition of the anti-ssrA oligonucleotide. Reactions were incubated at 37°C for 90 minutes for each round of selection, or from 7.5 to 90 minutes for the time course. At completion, reactions were placed at 4°C and diluted with 4 volumes (60 μL) of binding buffer (50 mM Tris-acetate (pH 7.5 at 4°C), 50 mM magnesium acetate, 150 mM NaCl, 1% Tween® 20, and 0.0 to 5.0% bovine serum albumin (BSA) or 0 to 20 mg/mL heparin). Meanwhile, for each reaction, 10 μL packed gel volumes of magnetic beads with selective markers or antibodies were washed three times with 50 μL bead wash buffer (50 mM Tris-acetate (pH 7.5 at 4°C), 50 mM magnesium acetate, 150 mM NaCl). Diluted iSAT reactions were added to washed beads and incubated at 4°C for 1 h with gentle rotation to suspend beads in solution. Reactions were then washed five or ten times with wash buffer (50 mM Tris-acetate (pH 7.5 at 4°C), 50 mM magnesium acetate, 150 to 1000 mM NaCl, and 0.05 to 5% Tween® 20), with 5 or 15 min incubations of each wash step at 4°C. Wash buffer was removed from the beads,50 μL elution buffer (50 mM Tris-acetate (pH 7.5 at 4°C), 150 mM NaCl, 50 mM EDTA (Ambion)) was added, and the beads were incubated at 4°C for 30 min with gentle rotation. Elution buffer was recovered from the beads, and rRNA was purified using a Qiagen RNEasy MinElute Cleanup Kit for rRNA analysis and/or RT-PCR.

### Reverse transcription polymerase chain reaction (RT-PCR)

For quantitative RT-PCR of 23S rRNA, RNA recovered from ribosome display was diluted 1:100 with nuclease-free water to dilute EDTA in the elution buffer. Diluted samples were used with the iTaq™ Universal SYBR® Green One-Step Kit (Bio-Rad) in 10 μL reactions following product literature. For quantitation, primers were designed for amplification of 23S rRNA using Primer3 software (**Supplementary Table S2**)^31^. Reactions were monitored for fluorescence in a CFX96™ Real-Time PCR Detection System (Bio-Rad). A standard curve was generated from a dilution series of 70S *E. coli* ribosomes (NEB) to ensure linearity of the assay (**Supplementary Figure S1**).

For recovery of rRNA from ribosome display, rRNA from capture ribosomes was purified with the RNeasy MinElute Cleanup Kit (Qiagen) and eluted with 14 μL nuclease-free water. Purified rRNA was used with the SuperScript® III One-Step RT-PCR System with Platinum® *Taq* DNA Polymerase (Invitrogen™) in 30 μL reactions following product literature. Primers were designed for use in both 23S rRNA recovery and *in vitro* operon plasmid assembly (see below) (**Supplementary Table S2**).

### *In vitro* operon plasmid assembly

For library construction or rRNA recovery, PCR was performed using Q5 polymerase with primers that amplify the bases 1962 to 2575 of the 23S rRNA gene. The primers also included unique cut sites for the off-site type lls restriction enzyme SapI, and the approximately 660 bp fragments and pT7rrnBΔ660 were digested with SapI in 1x CutSmart™ buffer (NEB) for 2 h at 37°C. DNA was purified, and inserts and plasmid were ligated at a 1:1 molar ratio with Quick Ligase (NEB) for 20 min at room temperature. The resulting plasmids were purified with DNA Clean & Concentrator-5™ (Zymo Research), eluted with nuclease-free water, and analyzed by NanoDrop to determine concentration.

### Isolation of individual sequence variants from RISE pools

Upon completion of RISE, reassembled rDNA operon pools were transformed into electro-competent POP cells for repression of the T7 promoter to prevent lethality of particular rDNA operon variants. Individual colonies were screened by colony PCR and sequencing. Colonies resulting in unclear sequencing reads were discarded. From the sequenced colonies, variants of interest were grown for plasmid preparation, and the purified plasmids were used in iSAT reactions as described above. Libraries, RISE pools, and isolated plasmids were submitted to the Genomics Core Facility at Northwestern University for sequencing.

### Deep sequencing of RISE selection cDNA pools

Mixed populations or clonal samples were prepared for deep sequencing beginning with a PCR of a 205 bp region containing the 6 randomized bases of the CR library near the center of the amplicon. These amplicons were processed for deep sequencing using NEBNext Ultra II FS DNA Library Prep Kit for Illumina (E7805S) and indexed using NEBNext Multiplex Oligos Index Primers Sets 1 and 2 (E7335S, E7500S). Samples were pooled and sequenced on an Illumina MiSeq v2 Micro flowcell with 2×150bp paired-end reads. Upon completion, .fastq files were trimmed using *sickle* (https://github.com/najoshi/sickle), filtered and aligned in Geneious, and the frequency of each genotype was counted and analyzed using a Python script. Only genotypes which were present with an absolute read count of 5 or greater in all sequenced pools were included in downstream analysis to lessen issues of increased error associated with observations at the bottom of the deep sequencing detection limit. This reduced the error in the dataset and restricted analysis to only those genotypes that were at 1.3% their initial frequency in the population or greater at the end of the selection, excluding most of the completely nonfunctional genotypes from correlation analysis.

### Squires strain analysis

Isolated variants were cloned into the appropriate *in vivo* vectors. Particularly, the T7 promoter was exchanged with the *P*_*L*_ promoter using standard molecular biology techniques (PCR, digestion, and ligation). Between 150-200 ng of sequence-verified plasmids were transformed into 50 μL of electrocompetent *E. coli* SQ171 cells, which lack chromosomal copies of the rDNA operon^22^. The transformants were recovered in 800 μL of SOC for 2 h at 37 °C. 1.2 mL of SOC supplemented with 100 μg/mL carbenicillin was added to the recovering culture, and the culture was further recovered overnight (16 h) at 37 °C. After the overnight recovery, 1 μL of culture was freshly inoculated into 100 μL of LB media containing 100 μg/mL carbenicillin in a 96-well plate with linear shaking maintained at 37 °C. Growth was monitored via absorbance at 600 nm, and at exponential phase (OD_600_ between 0.4 and 0.6), cultures were serially diluted to 10^−3^ to 10^−6^ OD_600_ and 2 μL of the dilution were spot plated onto agar plates containing 100 μg/ml carbenicillin, 350 μg/ml clindamycin, and/or 5% w/v sucrose.

### Estimating epistasis in the clindamycin-resistance selection

The epistasis value for each mutant was calculated using fitness values after the first round of selection in the presence of 500 μM clindamycin according to the following equation:

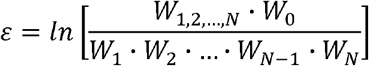

where *W*_l,2,…,*N*_ represents the fitness of the multi-mutant (measured as enrichment after one round of selection) that is a Hamming distance of *N* away from wild-type; *W*_0_ represents the fitness of wild-type; and *W*_l_, *W*_2_, and beyond represent the fitness of the individual point mutants that make up the multi-mutant. Thus, the numerator represents the observed fitness of the genotype, and the denominator represents its expected fitness, assuming each mutation contributes independently to the fitness of the genotype they comprise. The natural log of this fraction was then taken to obtain the epistasis value ε.

## ACKNOWLEDGMENTS

The authors would like to thank Adam Hockenberry for thoughtful review of the manuscript, and Jessica Stark for insightful comments on analysis. We also are thankful to Andreas Plückthun for use of the pRDV vector.

## AUTHOR CONTRIBUTIONS

M.J.H, B.R.F., and M.C.J. conceived this study. M.J.H. and B.R.F., D.J.Y. D.S.K., and E.D.C designed and executed the experiments. M.J.H., B.R.F., and M.C.J. wrote the manuscript and D.S.K. and D.J.Y. edited the manuscript. M.C.J. provided a supervisory role.

