## Supplementary Material for "*In vitro* ribosome synthesis and evolution through ribosome display"

^6^ Present address: Department of Chemical Engineering, Stanford University, Stanford, CA 94305, USA

^†^ Equal Contributions

Submitted to *Nature Communications*

**Supplementary Information**

**Supplementary Tables**

**Supplementary Table S1. Primers for plasmid constructions.** Bold letters indicate cut sites used for digestion and ligation, where an insert was generated from PCR with the indicated primers, and the resulting PCR product and backbone plasmid were digested with the same restriction enzymes prior to ligation. Gibson assembly was performed according to literature (Gibson *et al.*, 2009). ‘/Phos/’ indicates the use 5’ phosphorylation of primers or PCR product with the enzyme polynucleotide kinase (PNK) prior to ligation.

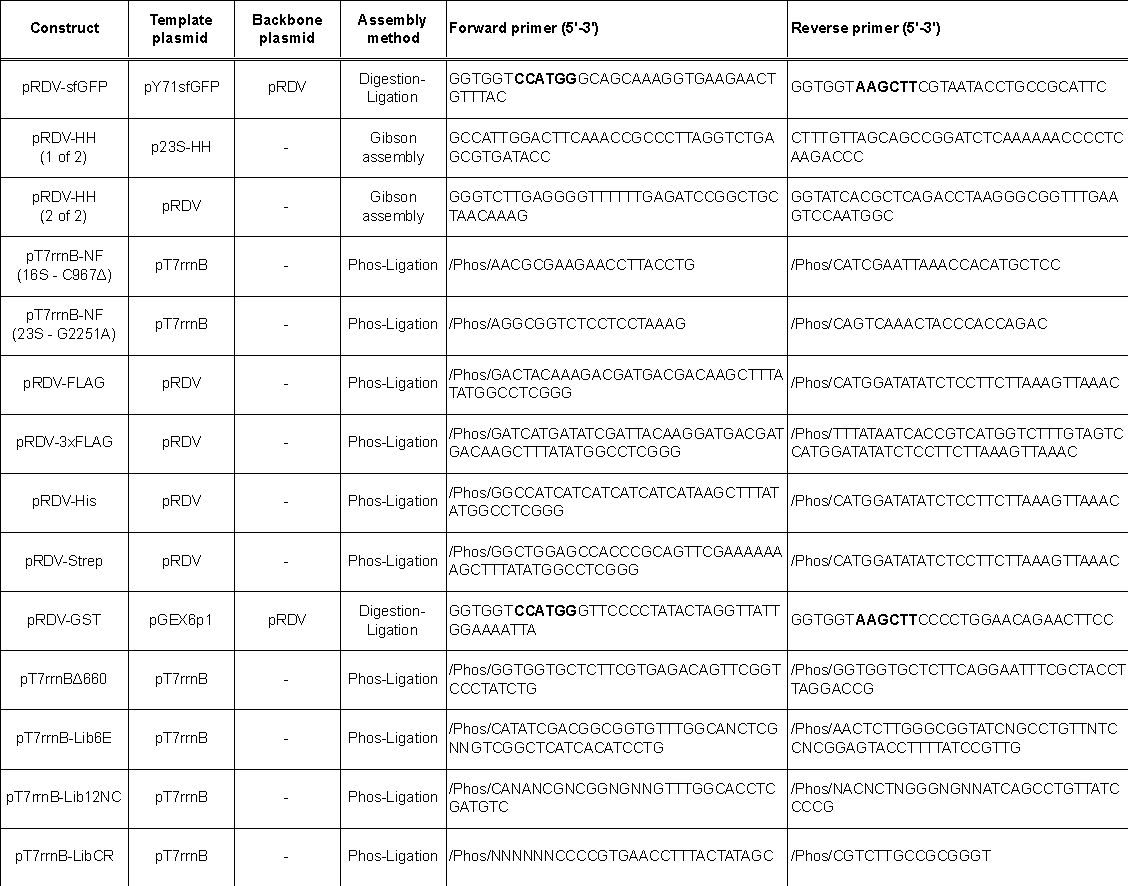

**Supplementary Table S2. Primers for RT-qPCR and RT-PCR of 23S rRNA.** Bold letters indicate cut sites used to reassemble operon from recovered and digested 23S cDNA and digested pT7rrnBΔ660.

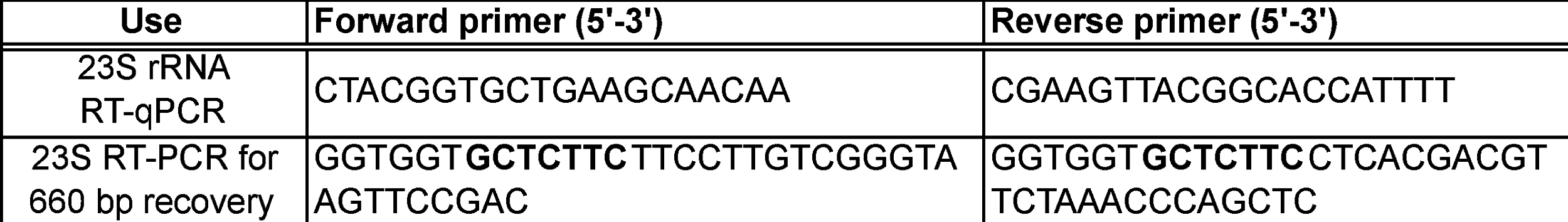

**Supplementary Table S3. Optimization of NaCl and Tween-20 concentration in wash buffer for RISE method with 3xFLAG-tag.** Values represent average specific capture of functional ribosomes relative to nonfunctional ribosomes as determined by RT-qPCR for two independent pairs of reactions (n=2).

| **Specificity using various wash buffers** | | **Tween-20 (%)** | | | |
| --- | --- | --- | --- | --- | --- |
|  |  | **0.05** | **0.25** | **1.00** | **5.00** |
| **NaCl (M)** | **0.15** | 126 | 150 | 129 | 119 |
|  | **0.30** | 118 | 110 | 87 | 89 |
|  | **0.50** | 92 | 93 | 101 | 85 |
|  | **1.00** | 85 | 94 | 71 | 84 |

**Supplementary Figures**

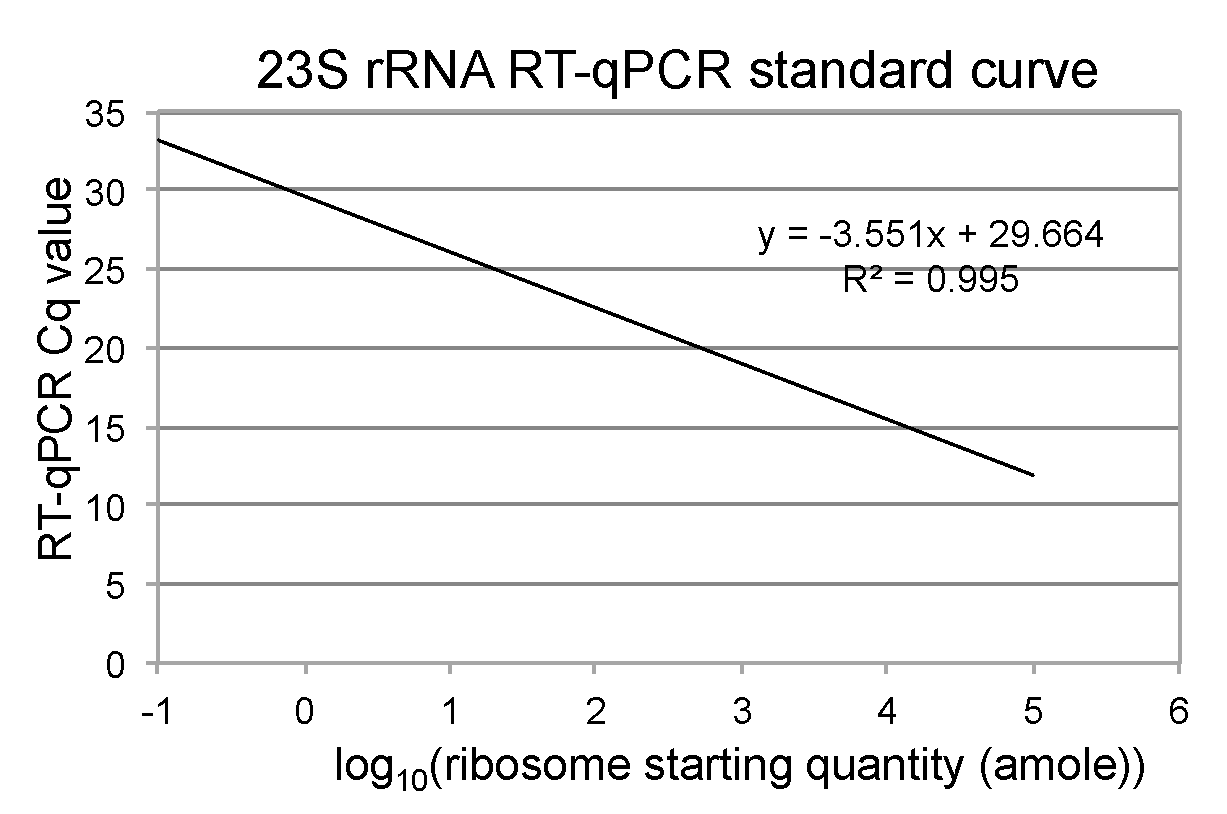

**Supplementary Figure S1. 23S rRNA RT-qPCR standard curve for dilution series of purified 70S *E. coli* ribosomes.** Purified 70S ribosomes were serially diluted 10-fold in nuclease-free water and used as templates in RT-qPCR.

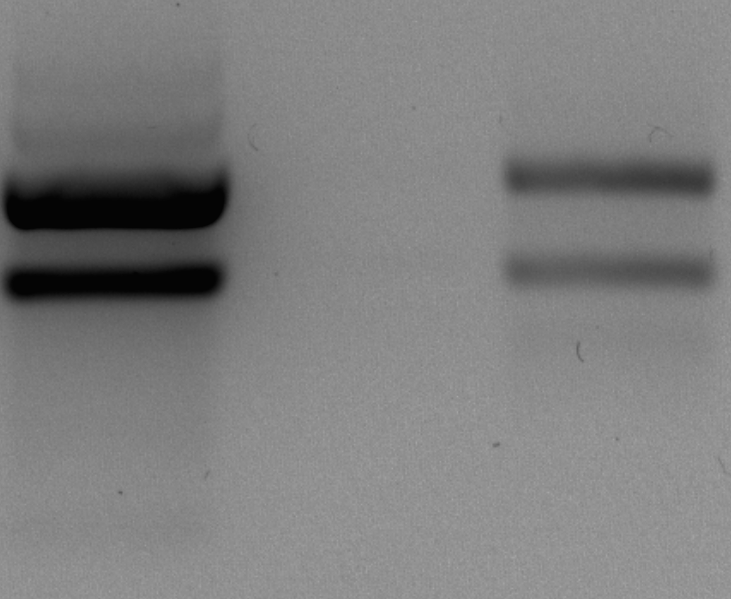

**Supplementary Figure S2. Ribosome capture by ribosome display.** Agarose gel shows ribosomes captured from iSAT reactions with nonfunctional (NF) or wild-type (WT) rDNA operon incubated for 1.5 h at 37°C. 4.5 pmol purified 70S ribosomes were run for comparison to represent the maximum theoretical number of iSAT ribosomes for 300 nM ribosomal proteins in a 15 μL reaction. Top band represents 23S rRNA and bottom band represents 16S rRNA. Gel representative of n=3 independent experiments.

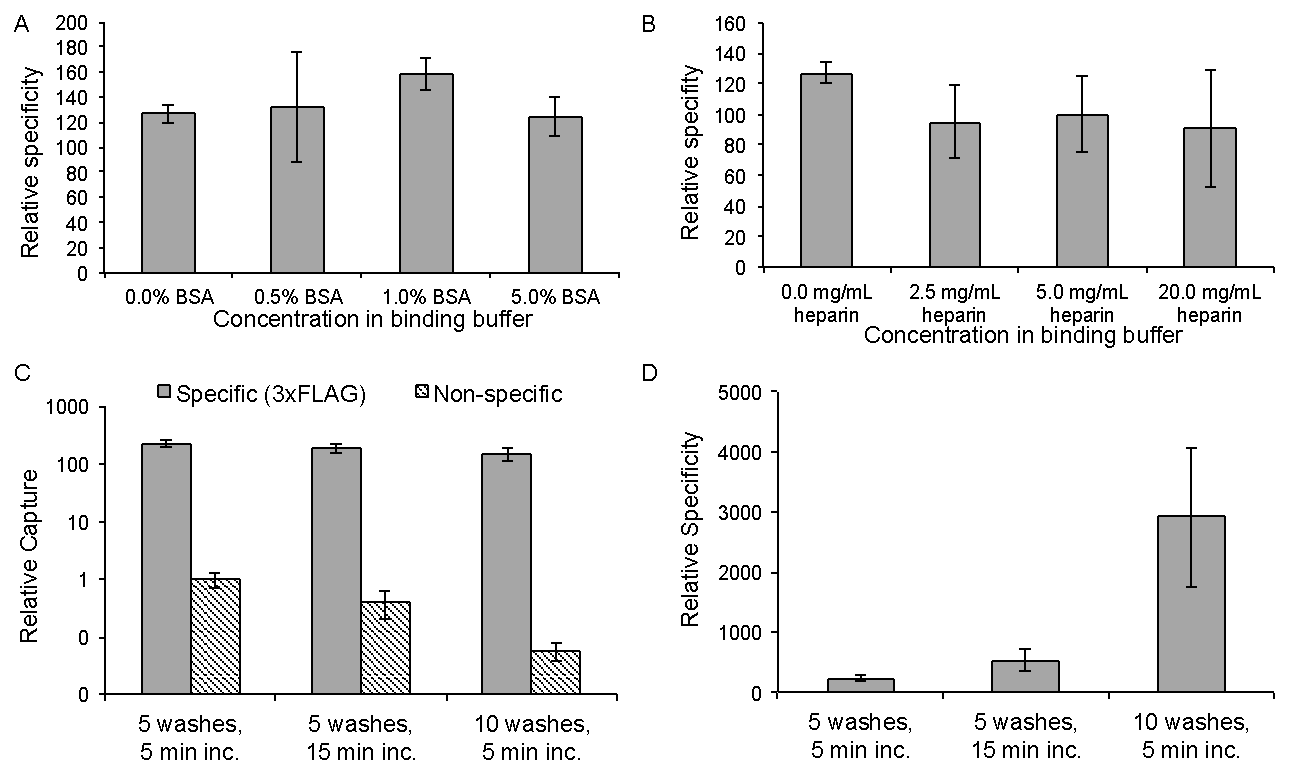

**Supplementary Figure S3. Optimization of binding and wash buffers for RISE using 3xFLAG-tag.** For the binding buffer, additives **(A)** BSA (% w/v) and **(B)** heparin were tested. For wash conditions, the number of washes and incubation of each wash were varied: **(C)** relative capture of iSAT ribosomes from specific capture (functional ribosomes) and non-specific capture (nonfunctional ribosomes) and **(D)** relative specificity of each wash condition. Values represent average specific capture of functional ribosomes relative to nonfunctional ribosomes as determined by RT-qPCR for three independent pairs of reactions (n=3). Error bars represent one s.d.

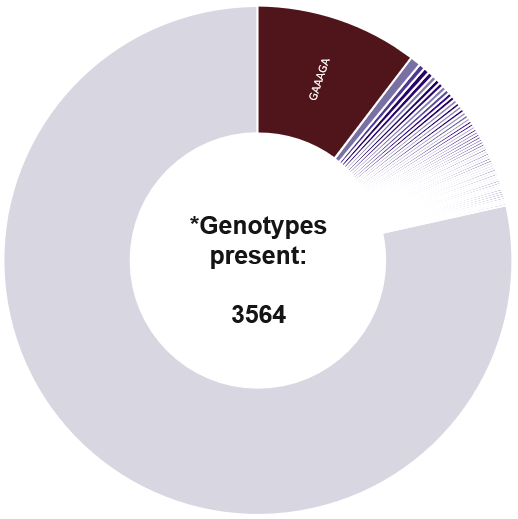

**Supplementary Figure S4. Initial genotype diversity in degenerate 6N CR library.** Sunburst plot depicting the initial genotype diversity in the CR library. Each slice represents the proportion of the initial population made up by the given genotype. The large gray slice represents all the genotypes at too low a frequency to depict graphically with an individual slice pooled into one group. The maroon slice indicates wild-type, revealing that the initial library was biased toward wild-type, likely due to incomplete cleavage of the wild-type plasmid when the library was built.

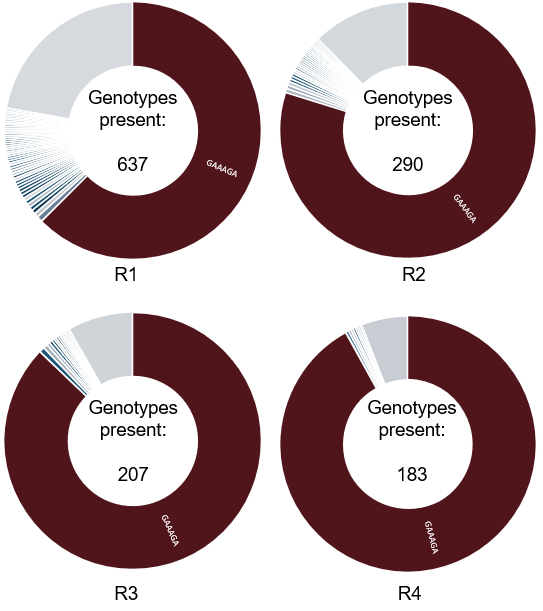

**Supplementary Figure S5. Genotype diversity across four rounds of selection on the CR library in 0 µM clindamycin.** While other genotypes are also seen increasing in frequency throughout the course of the selection, wild-type rapidly dominates the population over four rounds of selection in the absence of clindamycin. R‘X’ = Round X (e.g., 1, 2, 3, and 4).

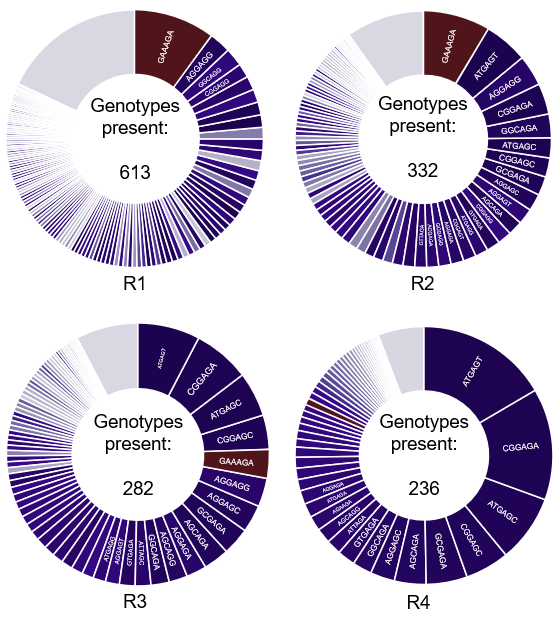
 **Supplementary Figure S6. Genotype diversity across four rounds of selection on the CR library in 500 µM clindamycin.**  Though wild-type starts out at high frequency, it is rapidly displaced by many clindamycin-resistant genotypes, indicating that our selection platform is specific for successfully translating ribosomes and is capable of selecting against wild-type under conditions where it does not translate efficiently. R‘X’ = Round X (e.g., 1, 2, 3, and 4).

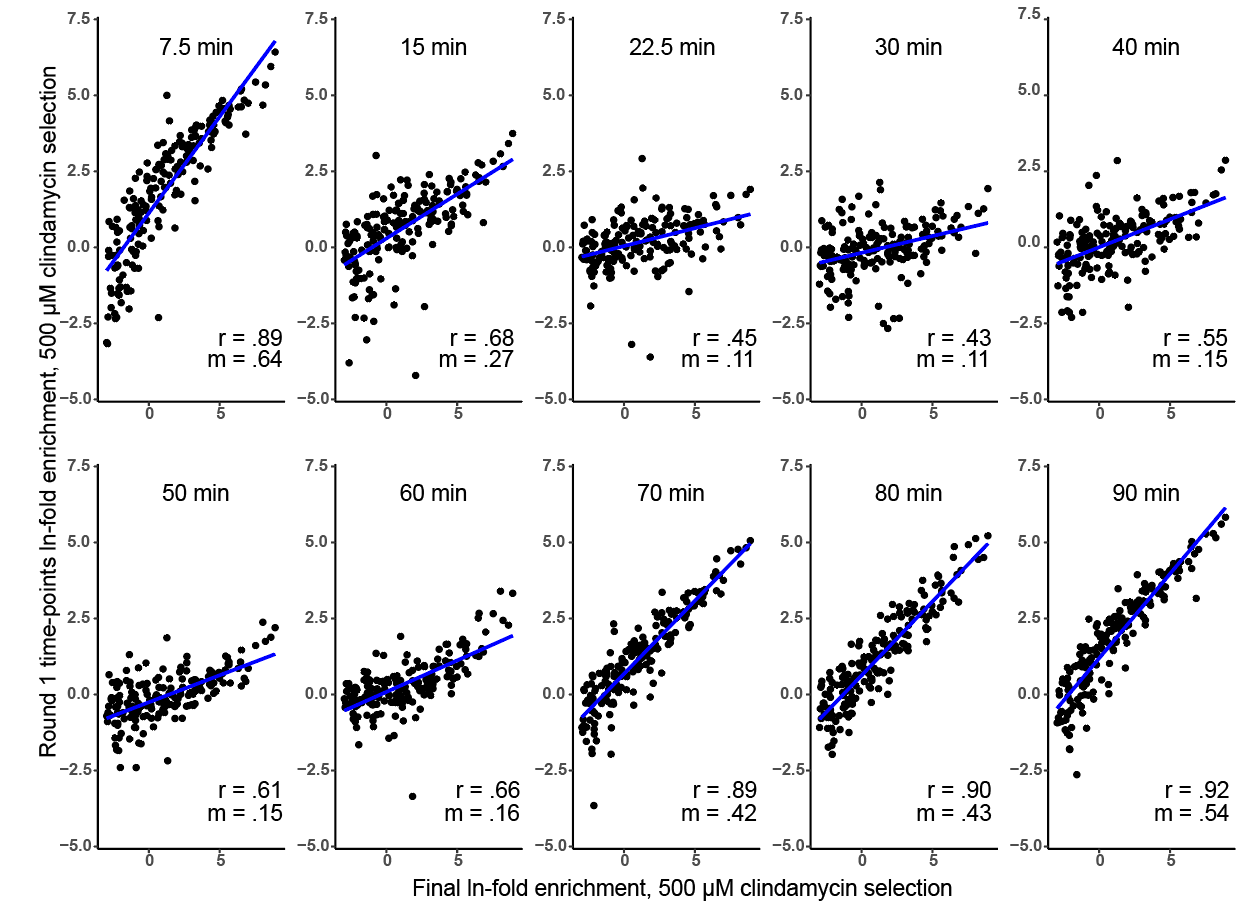

**Supplementary Figure S7. Time course of CR genotype enrichment from degenerate library in 500 µM clindamycin.** Each point represents a genotype whose ln-fold enrichment after four rounds of selection (x-axis) is plotted against its enrichment when the first round of selection is harvested at time points from 7.5 to 90 minutes. Since the most-enriched genotypes from the last round of selection are known to be highly active and clindamycin-resistant, the slope (m) and Pearson’s correlation coefficient (r) are measures of the strength and fidelity of the selection for active CR ribosomes at each time point, respectively. As expected, ribosome display reactions harvested at 70-90 minutes show efficient selection, in addition to the somewhat more surprising outcome of efficient selection for active ribosomes at the 7.5 minute time point.

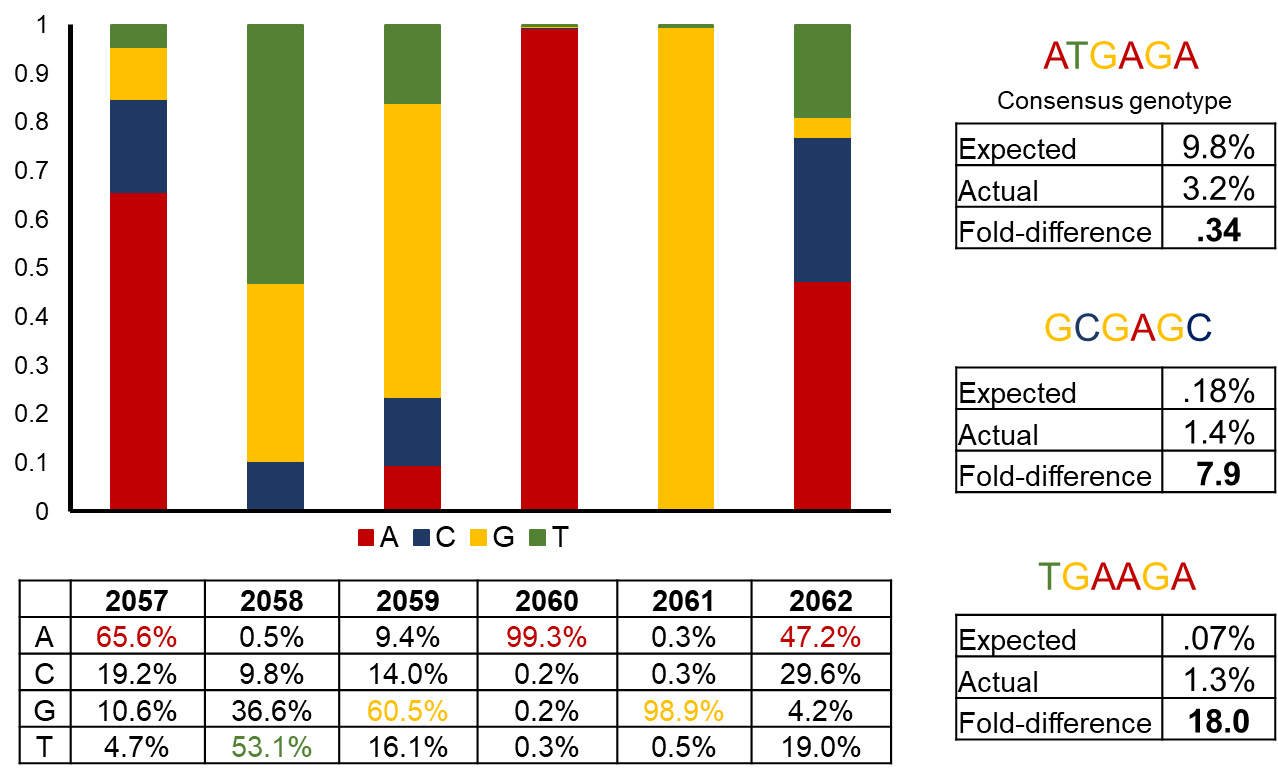
 **Supplementary Figure S8. Final nucleotide frequencies after four rounds of selection on 500 µM clindamycin.** While two positions (2060, 2061) have nearly converged after four rounds of selection, the remaining four show biases but retain significant diversity. The consensus genotype after four rounds is ATGAGA, though this genotype is present at approximately 3-fold lower concentration than expected by simply multiplying the frequency of each constituent base of the genotype together. Two other genotypes, GCGAGC and TGAAGA, however, are present at 7.9- and 18.0-fold higher frequency than expected. These genotypes would likely never be discovered by studying the effect of mutations at single positions on activity and clindamycin-resistance, highlighting the power of an evolutionary approach for discovering new ribosome variants.

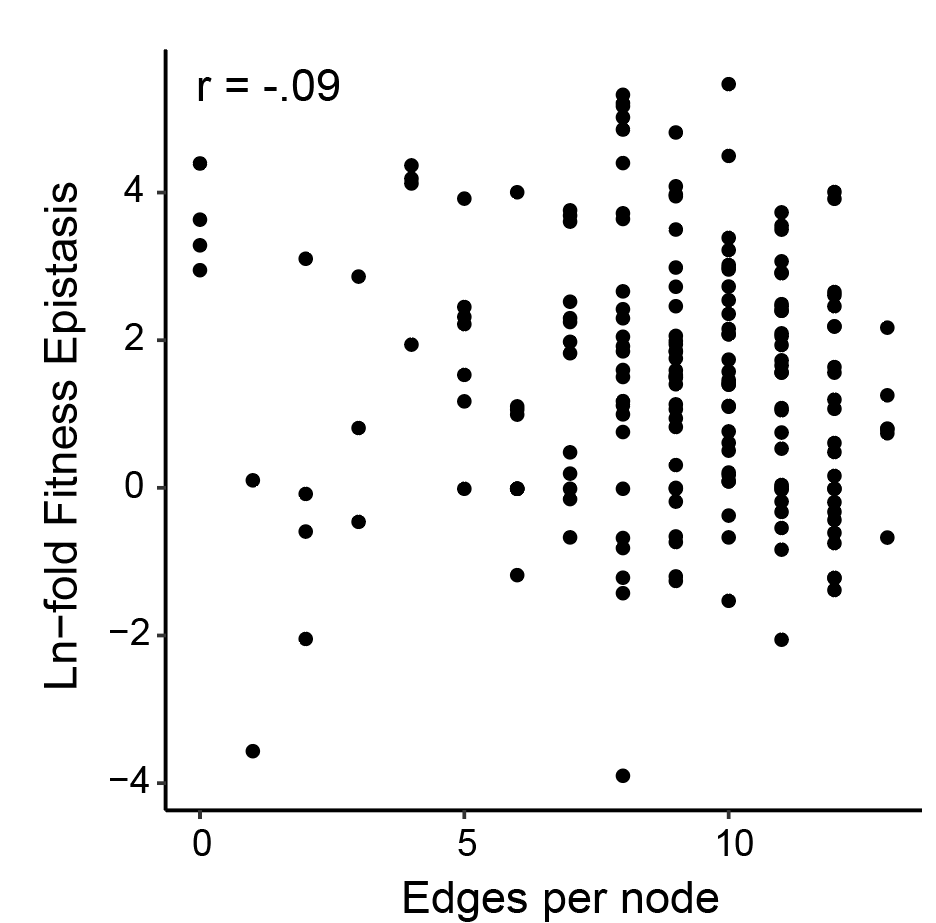

**Supplementary Figure S9. Epistasis values of clindamycin resistant genotypes plotted against edges per node.** Quantitative analysis of fitness and epistasis of the network of top Clindamycin Resistant genotypes shows that edges per node is not correlated with epistasis values.

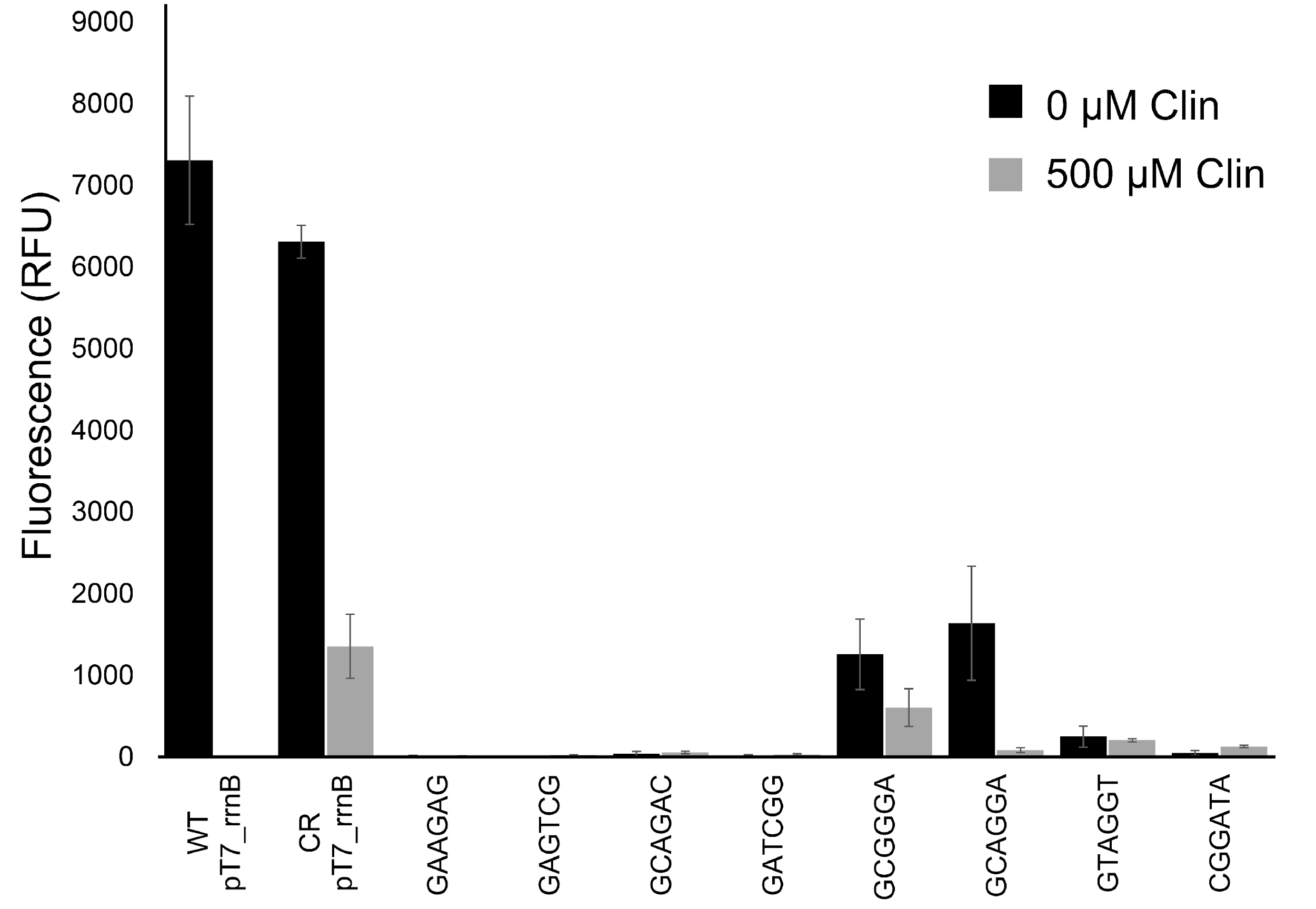

**Supplementary Figure S10. Function of extreme positive epistasis mutants in iSAT.**  While several genotypes displayed extreme positive epistasis values in our evolution experiment, the question remained whether these mutants retained any translation activity in iSAT or if their presence at the end of the selection was an artifact of selection. This is especially true for mutants differing from wild-type at positions 2060 and 2061, which are generally thought to be essential for ribosome function. A number of such mutants were cloned and tested for sfGFP production in iSAT. Several of these mutants (*e.g.,* GCGGGA, GCAGGA) retained measurable activity, indicating that they were indeed functional ribosomal mutants. Values represent mean values for four independent pairs of reactions (n=4). Error bars represent one s.d.
